# DNAm aging biomarkers are responsive: Insights from 51 longevity interventional studies in humans

**DOI:** 10.1101/2024.10.22.619522

**Authors:** Raghav Sehgal, Daniel Borrus, Jessica Kasamato, Jenel F. Armstrong, John Gonzalez, Yaroslav Markov, Ahana Priyanka, Ryan Smith, Natàlia Carreras, Varun B. Dwaraka, DNAm aging biomarkers community, community Longevity interventional studies, Albert Higgins-Chen

## Abstract

Aging biomarkers can potentially allow researchers to rapidly monitor the impact of an aging intervention, without the need for decade-spanning trials, by acting as surrogate endpoints. Prior to testing whether aging biomarkers may be useful as surrogate endpoints, it is first necessary to determine whether they are responsive to interventions that target aging. Epigenetic clocks are aging biomarkers based on DNA methylation with prognostic value for many aging outcomes. Many individual studies are beginning to explore whether epigenetic clocks are responsive to interventions. However, the diversity of both interventions and epigenetic clocks in different studies make them difficult to compare systematically. Here, we curate TranslAGE-Response, a harmonized database of 51 public and private longitudinal interventional studies and calculate a consistent set of 16 prominent epigenetic clocks for each study, along with 95 other DNAm biomarkers that help explain changes in each clock. With this database, we discover patterns of responsiveness across a variety of interventions and DNAm biomarkers. For example, clocks trained to predict mortality or pace of aging have the strongest response across all interventions and show consistent agreement with each other, pharmacological and lifestyle interventions drive the strongest response from DNAm biomarkers, and study population and study duration are key factors in driving responsiveness of DNAm biomarkers in an intervention. Some classes of interventions such as TNF-alpha inhibitors have strong, consistent effects across multiple studies, while others such as senolytic drugs have inconsistent effects. Clocks with multiple sub-scores (i.e. “explainable clocks”) provide specificity and greater mechanistic insight into responsiveness of interventions than single-score clocks. Our work can help the geroscience field design future clinical trials, by guiding the choice of interventions, specific subsets of epigenetic clocks to minimize multiple testing, study duration, study population, and sample size, with the eventual aim of determining whether epigenetic clocks can be used as surrogate endpoints.

## Introduction

The geroscience hypothesis proposes that targeting the underlying biological processes of aging may simultaneously prevent, delay, or mitigate multiple chronic diseases.^1^ Given the length of human lifespan, reliable aging biomarkers are essential for testing the efficacy of interventions.^2^ In this context, surrogate biomarkers that can predict long-term outcomes based on short-term measurements are particularly valuable as they can greatly speed up testing of new interventions.^3^ Additionally, discovery biomarkers, which reveal underlying biological pathways and mechanisms, play a critical role in identifying novel targets for intervention or prioritizing existing interventions for testing. ^2^ Together, these biomarkers will be critical in evaluating the potential of therapies to modulate aging and its associated diseases, and ultimately testing the geroscience hypothesis.

Aging biomarkers can be derived from diverse data sources, including clinical laboratory tests, functional phenotype evaluations, and molecular omics data.^2^ Each data type may offer unique and complementary insights into aging-related interventions. "Epigenetic clocks," based on DNA methylation (DNAm) at cytosine-guanine dinucleotides (CpGs), are among the most extensively studied biomarkers of aging.^4–6^ Numerous studies, including meta-analyses, have demonstrated that variations in epigenetic age among individuals of the same chronological age, in other words epigenetic age deviations or age residualized epigenetic ages, are strongly predictive of morbidity, mortality, and other age-related phenotypes. ^7–11^ Consistent sets of CpGs are measured by a given DNA methylation array, and many epigenetic clocks capturing different aspects of the aging process can be calculated from a single dataset. While there are some differences between different methylation arrays^12^, most clocks can be calculated from the widely available 450K, EPICv1, or EPICv2 data. Thus, even though epigenetic clocks are only one of many types of aging biomarkers, they are unique because they are well-suited to systematic comparisons across many studies and aging interventions.

Epigenetic clocks correlate with age and can serve as markers for aging processes. The first epigenetic clocks, such as Hannum^13^, Horvath Multi-Tissue (Horvath1) ^14^, and Horvath Skin and Blood (Horvath2) ^13,151513,15^, were trained to predict chronological age and are referred to as Generation 1 clocks. Subsequent clocks focused on measuring phenotypic aging, mortality risk, or the pace of aging. This category includes PhenoAge^9^, GrimAge^8^, and DunedinPoAm38^16^, collectively known as Generation 2+ clocks. It was later found that clocks suffered from significant technical noise that degraded their test-retest reliability.^17^ Thus, existing clocks were modified through two distinct approaches: using principal components of CpGs (e.g. PCPhenoAge or PCGrimAge), or by incorporating only CpGs with high test-retest reliability into the training paradigm (e.g. DunedinPACE^11^). Here, we refer to these as "Generation Reliable" clocks. Recently, acknowledging that aging is multifaceted and involves many biological processes, new clocks emerged to better explain the underlying mechanisms captured by previous generations. Whereas prior clocks report a single score for the whole body, these newer clocks typically involve a panel of multiple scores that can capture heterogeneity in aging. These clocks, which we refer to as "Generation Explainable," (Gen X) decompose the single-score or whole-body clocks into components to provide greater insight into the biological processes of aging. For example, SystemsAge^10^ provides 11 different scores aimed at quantifying aging in different physiological systems using only blood DNA methylation data. CausalAge disentangles damaging and adaptive changes as two separate scores.^18^ OMICmAge^19^ and related clocks utilize DNAm surrogates of proteins, metabolites, and clinical phenotypes. These advancements reflect a growing sophistication in the design and application of epigenetic clocks, enhancing their utility in aging research and potential clinical applications.

There are also a variety of DNAm biomarkers that predict smoking^8^, cell composition^20^, serum protein levels^8,21^, metabolite levels^19^ and other molecular and phenotypic features. Some, but not all, of these change with age. Many of these are components of epigenetic clocks and can help explain why they might change. To avoid confusion, and to be more precise, we use “DNAm biomarkers” as the umbrella term and reserve “epigenetic clocks” for those DNAm biomarkers specifically meant to capture aging.

Most validation of epigenetic clocks has focused on their ability to predict baseline disease status or future mortality and morbidity outcomes ^7^. By comparison, relatively little is known about the responsiveness of epigenetic clocks and other DNAm biomarkers to various interventions.^22^ Validating this responsiveness is an important step towards determining whether DNAm biomarkers may be surrogate endpoints in interventional clinical trials. It will be further necessary to determine whether the degree of change in biomarkers is predictive of change in aging outcomes, but such studies will be very expensive and lengthy, and therefore a roadmap of which biomarkers are responsive to which interventions will be useful in planning further trials.

A few recent studies have investigated DNAm biomarker responsiveness to individual interventions. The CALERIE trial, which focused on calorie restriction, calculated 11 DNAm biomarkers including DunedinPACE, GrimAge, PhenoAge, Horvath1, and Hannum, along with their principal component (PC) counterparts. Only DunedinPACE and PhenoAge showed significant decreases, with the latter result being unclear because the reliable equivalent PCPhenoAge did not show a significant change.^22,23^ In the DAMA trial of a plant-food-rich diet and exercise intervention, only GrimAge was calculated which showed a decrease of 0.66 years.^23,24^ The MDL study of a lifestyle intervention calculated only Horvath1 and found a decrease of 3.2 years.^25^ The D-SUNNY study, focusing on vitamin D supplementation, measured Horvath1 and Hannum, with a 1.86-year decrease seen only in Horvath.^26^ The Umbilical Cord Plasma Concentrate study measured several biomarkers, including GrimAge, PhenoAge, Horvath1, Horvath2, and Hannum. Among these, only GrimAge showed a reduction.^27^ The B12 Folate and MOFs dietary supplement studies calculated Horvath1, and observed a reduction in a subgroup but not in the overall cohort. ^28^ In the Senolytics DQF and DQ trials, multiple biomarkers such as DunedinPACE, PCGrimAge, PCPhenoAge, PCHorvath, and PCDNAmTL were calculated. In both trials, PCHannum showed an increase in epigenetic age, along with other biomarkers in the DQ trial.^29^

We have consolidated the results from various trials and studies in Figure 1A, which highlights several concerns that limit our ability to draw conclusions about both clocks and interventions from the literature. Most notably, different studies typically utilize different sets of epigenetic clocks, with some studies reporting many clocks and others reporting one or a handful of clocks. This inconsistency has two major consequences: first, this makes it very difficult to compare results from individual studies. For example, multiple studies examined only the original Horvath multi-tissue clock (Horvath1). For the two studies that only examined Horvath1 and found no change, it is possible other clocks would have detected a change, especially considering that CALERIE and the Umbilical Cord Plasma study also found no change in Horvath1 but did find changes in DunedinPACE and GrimAge respectively. Conversely, for the two studies that only examined Horvath1 and found decreases, it is plausible no other clocks would show a change, or another clock might increase, raising questions about the significance of the detected change. If only a single clock changes and no others, it is uncertain whether it is a false positive or if that clock is simply the most sensitive to interventions. The large number of clocks raises the possibility of publication bias where only positive results for specific combinations of clocks and interventions are reported. It is difficult to compare different studies of related interventions - for example the three dietary studies CALERIE, MDL and DAMA found changes in different sets of clocks (DunedinPACE, Horvath1, and GrimAge respectively), but CALERIE also calculated Horvath1 and GrimAge and found no change. It seems puzzling why related interventions might affect completely different clocks. A second major consequence of inconsistent clocks between studies is that we cannot draw any global conclusions about clocks and interventions. For example, we cannot determine if there is any epigenetic clock that is most responsive to multiple interventions, or if there is any intervention that changes epigenetic age consistently either across clocks or across studies of the same intervention. While it is possible that different clocks are impacted by different interventions, there is no way to currently assess that. In addition to clock choice, it is possible that systematic differences between studies stem from different sample sizes, durations, designs, and populations, but there is no harmonized set of study metadata to investigate this issue. All of these concerns make it very difficult for researchers designing future clinical trials to leverage existing literature to select interventions and clocks that they wish to test.

**Figure 1:**
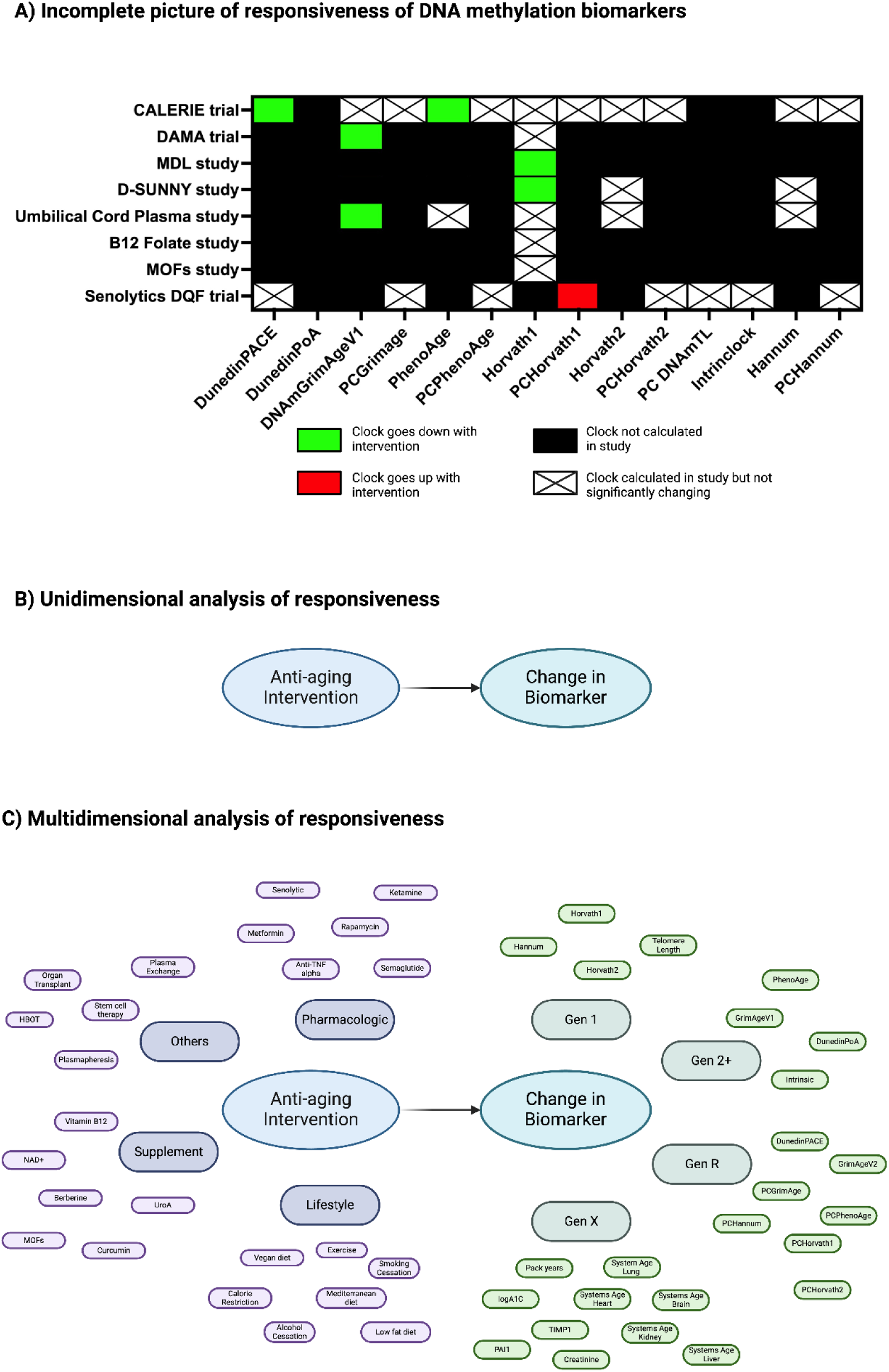
Current paradigms to study responsiveness of DNAm biomarkers and their limitations. A) Incomplete Picture of Responsiveness: A matrix summarizing existing literature illustrating the effects of various interventions on different DNA methylation biomarkers. After intervention, the clocks may increase (red), decrease (green), not change (cross), or not be calculated at all (black). . B) Unidimensional Analysis of Responsiveness: A traditional application of the concept of responsiveness to aging biomarkers may only consider the change in one biomarker after one intervention. C) Multidimensional Analysis of Responsiveness: Given that there are many aging interventions and many clocks, a complex framework is needed which categorizes anti-aging interventions and biomarkers and underscores the diverse influences of different anti-aging intervention types on various generations of DNA methylation biomarkers.

We set out to test the hypothesis that “Epigenetic aging biomarkers are responsive to aging interventions”. We reasoned that the diversity of both aging interventions and epigenetic clocks necessitates an integrative analysis that systematically explores the interactions between all interventions and the entire spectrum of DNA methylation biomarkers. We present this integrative analysis as an alternative approach to the existing fragmented literature where studies report the impact of a single intervention on limited lists of DNAm biomarkers that are discordant between studies. Thus, to test our hypothesis systematically, we have curated and harmonized 51 anti-aging studies with DNA methylation data, standardized metadata, and calculated a consistent set of 16 epigenetic clocks and Y other DNAm biomarkers. Results are compared for the first time across interventions, revealing which biomarkers and interventions show the strongest and most consistent responses.

## Results

### Data curation and analysis pipeline

Curating and analyzing a large database of interventions required a robust and standardized analysis pipeline. For this purpose we built a novel pipeline that combines standard practices and packages used in aging biomarker analysis (Figure 2). First, we created a list of publicly (GEO and EMBL) and privately (TruDiagnostic) available longevity interventional clinical trials and studies. For each study we curated study level metadata such as intervention type, study population, diseases in population (if any), age range, and percent female. Second, we harmonized metadata in each study to have standardized columns such as: Age, Sex, Sample ID, Individual ID, Follow-up time from baseline draw and sample type (control vs subject). Third, we calculated over 110 DNAm biomarkers for each of these datasets using the MethylCIPHER package ^30^, updated to include more clocks. Fourth, we adjust for chronological age to obtain age residuals for all DNAm biomarkers in every dataset. Fifth, we normalized the biomarker age residuals in each dataset with the standard deviation of the age residual in the Health and

**Figure 2:**
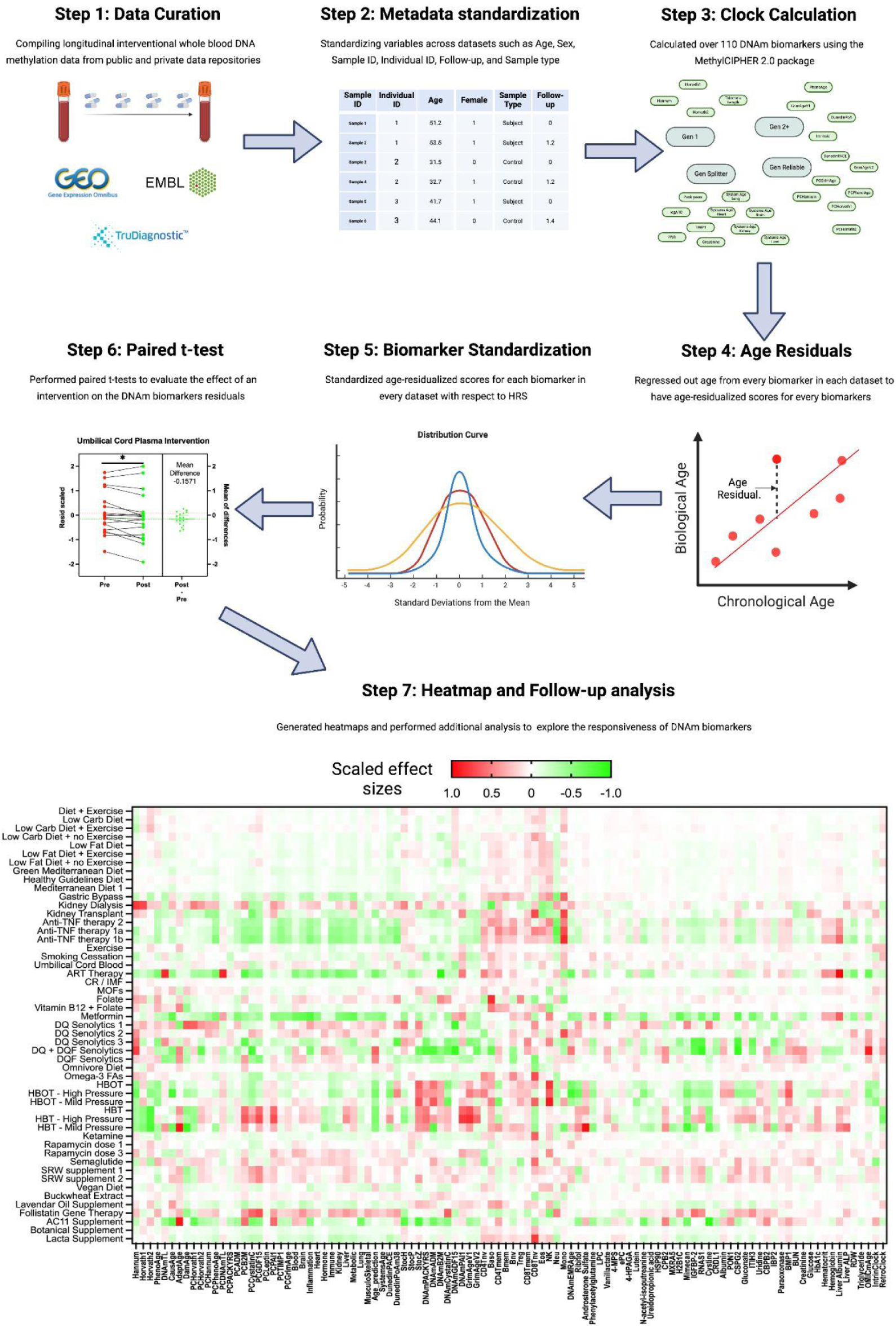
Standardized and comprehensive data curation and analysis pipeline for longitudinal interventional DNA methylation biomarker studies. Step 1) Data Curation: This initial step involved gathering a diverse array of whole blood DNA methylation datasets from multiple longevity-focused clinical trials and studies from various public (GEO and EMBL) and private (TruDiagnostic) repositories and standardizing their study level metadata. Step 2) Metadata Standardization: Metadata for each study was harmonized to ensure consistency across datasets. Standardized variables included Age, Sex, Sample ID, Individual ID, Follow-up time from baseline draw, and Sample type (categorizing control vs. subject). This standardization was crucial for enabling reliable comparisons and analyses. Step 3) Clock Calculation: Using the MethylCIPHER 2.0 package, over 110 DNAm biomarkers were calculated for each dataset. Step 4) Age Residuals calculation: Age was regressed out from every biomarker in each dataset to isolate age-residualized scores. Step 5) Biomarker Standardization Age-residualized scores: For each biomarker, in every dataset, age residualized scores were standardized relative to the Health and Retirement Study. This normalization was essential for ensuring that comparisons across different datasets were accurate and meaningful. Step 6) Paired t-test Analysis: Paired t-tests were performed on pre- and post-intervention samples to evaluate the effects of interventions on DNAm biomarkers. This statistical analysis enabled the determination of the significance of changes in biomarkers due to interventions. Step 7:) Heatmap and Follow-up Analysis to explore the responsiveness of DNAm biomarkers. The heatmap shows scaled effect sizes from red (positive effect) to green (negative effect), visualizing intervention impacts on biomarkers.

Retirement study to make it comparable across interventional studies ^31^. Sixth, we performed paired t-test analysis for pre and post intervention samples. Finally, we generated heatmaps of the mean scaled effect size and performed additional analyses to understand more about the responsiveness of these biomarkers. The significance threshold used in this analysis is set at 0.05, without corrections for multiple hypothesis testing. This approach is justified for several reasons: 1) Multiple testing corrections assume test independence, which is not applicable here since the clocks are highly intercorrelated, risking the exclusion of true positives. 2) Interventions of similar categories may show correlated effects, further complicating correction. 3) The ultimate aim is to simplify the process by identifying clocks that consistently capture intervention effects. This would allow us to eventually focus on a smaller, reliable set of DNAm biomarkers, either through individual selection or by developing ensemble biomarkers. By reducing redundancy, we minimize the need for multiple hypothesis testing across hundreds of biomarkers, leading to a more streamlined and meaningful analysis.

### Longevity interventions have varying degrees of effects on DNAm biomarkers

We first focused on 16 prominent epigenetic clocks that are not focused on any particular aspect of aging. These included Generation 1 biomarkers (Horvath1, Horvath2, and Hannum) Generation 2+ biomarkers (PhenoAge, GrimAgeV1, DunedinPoAm38), Generation Reliable clocks that are updates of earlier clocks (PCHorvath1, PCHorvath2, PCHannum, PCPhenoAge, PCGrimAge, GrimAgeV2, DunedinPACE), and Generation Reliable clocks optimized for reliability during initial development (SystemsAge, OMICmAge, DNAmEMRAge). We plotted the effect size of all 51 interventions for these 16 DNAm biomarkers (Fig 3A) with the non-significant effect sizes grayed out (p-value <0.05). Some interventions seemed to affect more DNAm biomarkers (positively or negatively) than others. To quantify this without making any initial assumptions about which biomarkers are more important, we plotted all effect sizes mapping to the 16 different biomarkers for an intervention and then performed a one way t-test on effect sizes of all interventions checking for deviations from 0 effect size (Fig 3B). 19 interventions significantly decreased epigenetic age across the 16 biomarkers, 5 significantly increased epigenetic age across the 16 biomarkers and the remaining 26 had no significant effects.

**Figure 3:**
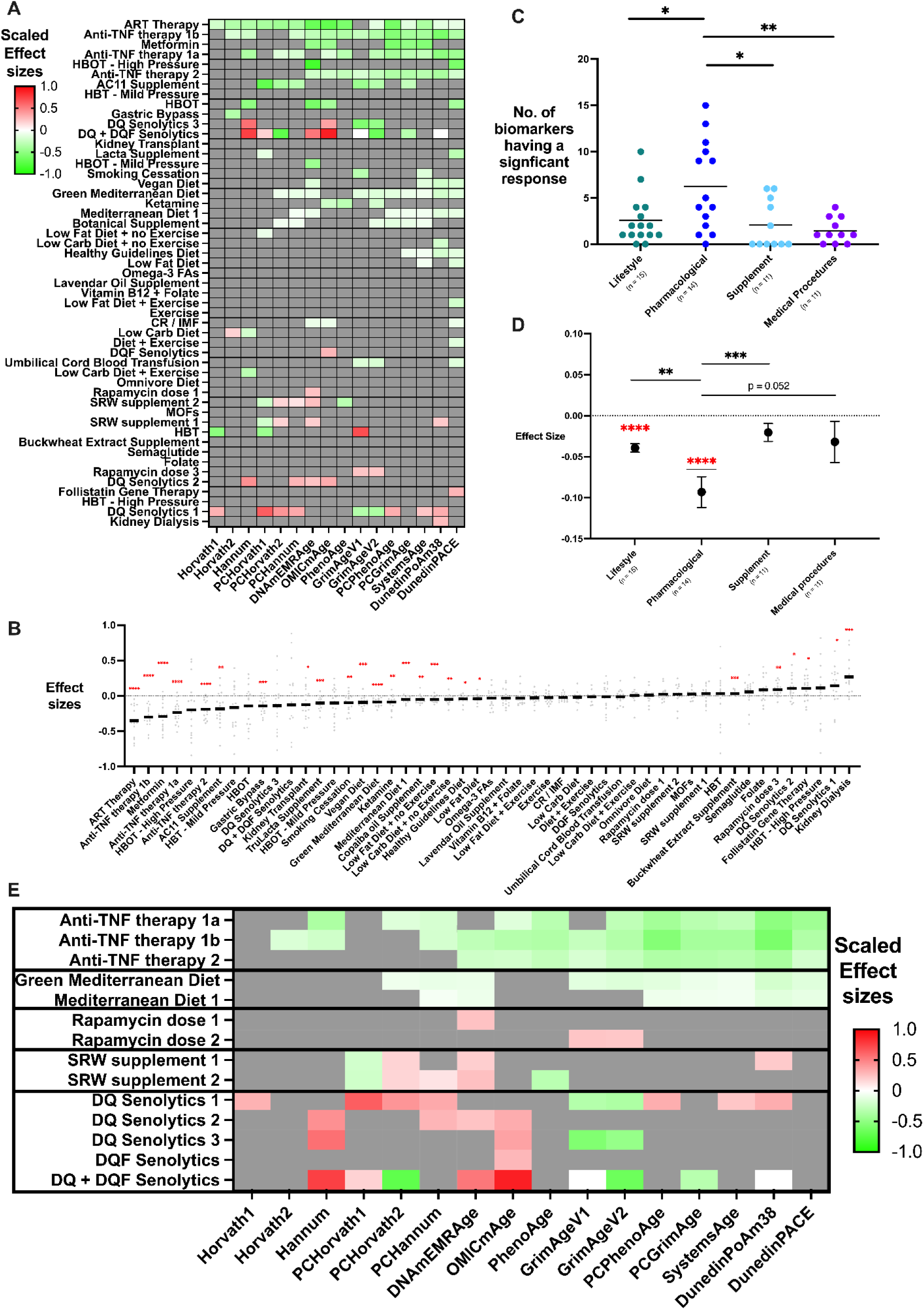
Interventions and their effects on 16 prominent DNAm biomarkers. A) Heatmap of biomarker responses to different interventions. The x-axis lists the biomarkers, while the y-axis lists the interventions. Red indicates increase in epigenetic age, green indicates decrease in epigenetic age, while gray denotes no significant effect. B) Categorical scatter plot comparing effect sizes of different intervention types across all 16 biomarkers. For each intervention, each point corresponds to one clock and solid lines indicate mean effect size across clocks. Asterisks indicate interventions that significantly alter epigenetic age averaged across all 16 DNAm biomarkers (* p < 0.05, ** p < 0.01, *** p < 0.001). C) Categorical scatter plot counting number of biomarkers with statistically significant response, regardless of direction, with studies separated into 4 categories (Lifestyle, Pharmacological, Supplements, and Medical Procedures). Each point corresponds to one interventional study. Significant differences between categories are indicated by asterisks (* p < 0.05, ** p < 0.01), compared to Pharmacological as the reference category.. D) Effect Size Comparison Across Intervention Categories: This plot compares effect sizes across different intervention categories. The y-axis represents the effect size, while the x-axis categorizes interventions into Lifestyle, Pharmacological, Supplements, and Medical Procedures. Significant differences between categories are indicated by asterisks (** p < 0.01, *** p < 0.001). E) Heatmap of interventions where translAGE contains multiple studies of the same intervention, allowing consistency between studies to be examined. Red indicates increase in epigenetic age, green indicates decrease in epigenetic age, while gray denotes no significant effect.

### Certain longevity intervention categories elicit stronger responses from DNAm biomarkers

Given the diversity of interventions we collated in our database, we asked whether some types or categories of longevity interventions have greater effects on DNAm biomarkers than others. We categorized interventions in our database into 4 categories: Lifestyle (Diets, exercise and more), Pharmacological (Metformin, Rapamycin, Semaglutide, Ketamine, Anti-TNF therapy and more), Supplements (Omega 3FA, Folate and more) and Medical Procedures (Hyperbaric Oxygen Therapy, Organ Transplant, Gene Therapy and more). We then performed two analyses. In the first analysis we simply counted the number of significant biomarkers (out of the 16 biomarkers previously mentioned) per intervention and then plotted the number of significant biomarkers on a scatter plot by intervention category (Fig 3C). Finally, we performed a paired t-test with every comparison, which revealed that only the number of significant biomarkers in the Pharmacological category were significantly higher than any other category. Pointing at the fact that potentially, pharmacological interventions might be eliciting a stronger response from DNAm biomarkers. To further test and quantify this hypothesis we performed another one-way t-test but this time on all clock effect sizes in an intervention category (Fig 3D). Only two category of interventions seem to have a significant effect on decreasing epigenetic age: Pharmacological (mean effect size = -0.09307, t-value = 4.948) and Lifestyle (mean effect size = -0.0393, t-value = 7.579) with Pharmacological having a larger effect size. Additionally, when t-tests were performed between different categories only Pharmacological seemed to have significant effect sizes compared to the other categories (Lifestyle: P-value = 0.0063; Supplement = 0.0009, Medical Procedures = 0.0517). Both these analyses point at the fact that Pharmacological interventions may be driving a larger effect from DNAm biomarkers than other intervention types.

### Consistency between biomarkers and studies supports *bona fide* effects of longevity interventions

Our compiled database includes a number of replicate studies of the same intervention, as well as multiple clocks that measure related constructs (e.g. multiple mortality clocks). We reasoned this would be useful for determining if an intervention is indeed modifying DNAm biomarkers in a consistent manner which would support it as an effective longevity intervention. While false positives for a given clock/intervention combination are likely to occur, it is unlikely that multiple clocks and multiple studies would repeatedly yield the same false positives. Thus we propose that interventions with *bona fide* effects of aging biomarkers should fulfill two rules: first, the intervention should modify DNAm biomarkers of a given generation to the same magnitude and direction in a particular study (Rule 1), and second study of the same intervention should modify the same biomarkers (Rule 2). In our database, we can identify interventions that abide by both rules, one of the rules or neither of the rules (Figure 3E). Anti-TNF Therapies in patients from multiple studies (Arthritis and IBD) modify nearly all Gen 2+ biomarkers (both reliable and otherwise) to very similar magnitude, fulfilling both rules and supporting anti-TNF therapies as a potential method to prevent pathological aging in individuals with autoimmune disorders. Similarly, two different types of Mediterranean diets, performed in healthy cohorts, decrease similar subsets of Gen 2+ DNAm biomarkers, satisfying both Rule 1 and Rule 2. On the other hand, we see an example of two SRW supplement studies which change the same biomarkers (satisfying Rule 2), but there is no consistent change in direction across those biomarkers (not satisfying Rule 1 - PCHorvath1 and PhenoAge decrease while PCHorvath2, PCHannum and DNAmEMRAge increase). This inconsistency between biomarkers generates uncertainty about the effects of such an intervention on aging. Similarly, the five different Senolytics studies have multiple biomarkers changing in different direction within a study (GrimAgeV1 and V2 vs others, Not Satisfying Rule 1) and sometimes the same biomarker changing in opposite direction across studies (PCHorvath2, Not Satisfying Rule 2). This example suggests that the interventional effects of senolytics, at least from an epigenetic aging perspective, may be inconsistent.

### Reliable Gen 2+ biomarkers demonstrate the highest responsiveness

In addition to comparing interventions, we aimed to compare the responsiveness of the 16 DNAm biomarkers discussed previously. As an initial survey, we counted the number of interventions for which each DNAm biomarker decreased or increased significantly and compared these on a scatter plot (Figure 4A). Most Gen 2+ reliable biomarkers such as SystemsAge, GrimAgeV2, PCGrimAge, PCPhenoAge, and DunedinPACE, decreased significantly in multiple interventions and increased in none or few interventions. Notably, the only Gen 2+ biomarkers that increased in more than one study were GrimAgeV1 and DunedinPoAm38, which supports utilizing their more reliable counterparts instead. Meanwhile, most Gen 1 biomarkers, such as Horvath1, Horvath2, and Hannum, sporadically increased or decreased in a handful of different interventions, lacking any clear trend. DNAmEMRAge and OMICmAge changed in many interventions, though interestingly they were a mix of increases and decreases, which may reflect the complexity of the multi-omic and phenotypic data on which they were trained. Among all the biomarkers, DunedinPACE was the most responsive to a deflection in biological age, significantly decreasing in 16 interventions and increasing in only 1. To further quantify responsiveness, we performed a second analysis where we plotted all the effect sizes for a single clock from all interventions and performed a one-way t-test checking for deviations from zero for each clock. We reasoned that since all interventions in our dataset are targeted at ameliorating pathological aging potentially by reducing biological age, then if the DNAm biomarkers are indeed responsive they should show a significant decrease when all effect sizes were pooled together. This analysis revealed that all Gen 2+ Reliable biomarkers showed significant decreases. These include GrimAgeV2 (mean = -0.0814, p= 0.0075), PCPhenoAge (mean = -0.08876, p = 0.003), PCGrimAge (mean = -0.07843, p =0.0003), SystemsAge (mean = -0.05859, p = 0.0043) and DunedinPACE which had the largest decrease (mean = -0.0891, p = 0.0012). In contrast, only a single Gen 1 (reliable) clock significantly decreased across all interventions (PCHorvath1 mean = -0.06674, P-value = 0.029), and a single Gen 2 non-reliable DNAm biomarker decreased (PhenoAge mean = -0.06735, p = 0.0408). Overall, both analyses revealed the same insight: reliable Gen 2+ biomarkers are responsive to interventions, with DunedinPACE showing the strongest response. This suggests potentially two criteria for a biomarker to be responsive 1) it needs to be reliable 2) it needs to be a mortality or rate of aging predictor.

**Figure 4:**
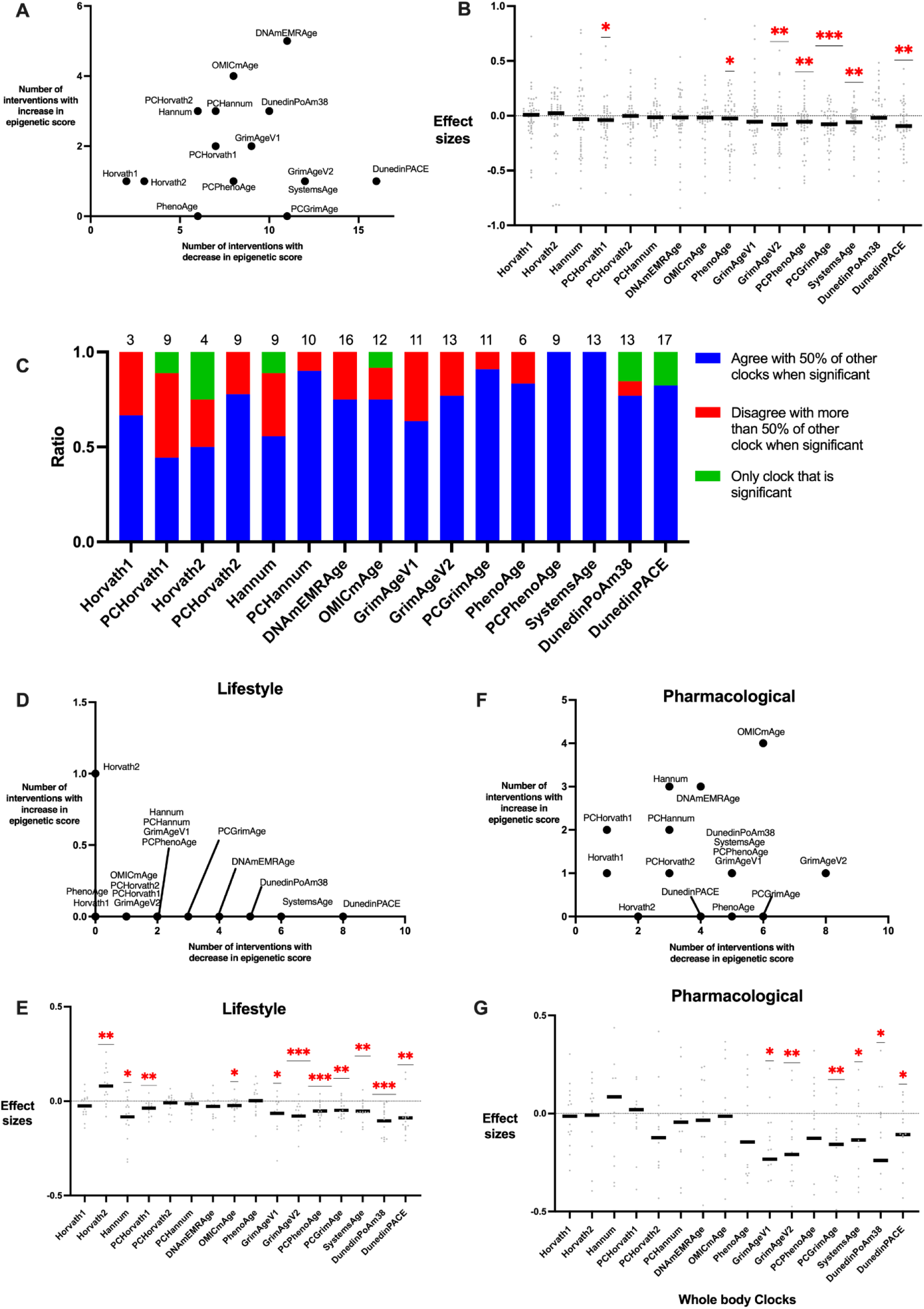
Response of 16 prominent DNAm Biomarkers to Interventions. A) Scatter plot of count of interventions where each DNAm biomarker decreases (X-axis) or increases (Y-axis) B) Categorical scatter plot of effect sizes of interventions on DNAm biomarkers where each point is one intervention study: Significant effect sizes are marked with red asterisks (* p < 0.05, ** p < 0.01, *** p < 0.001), while solid lines indicate the mean effect size. C) Bar plot of agreement and disagreement among DNAm biomarkers. For each clock, the number of interventions leading to significant changes were counted (denoted by the numbers at the top). Then we categorized those significant changes into agreeing with over 50% of other clocks showing a change (blue), disagreeing with over 50% (red), or being the only significant clock change (green). D) Scatter plot of count of lifestyle interventions where each DNAm biomarker decreases (X-axis) or increases (Y-axis) E) Categorical scatter plot of effect sizes of lifestyle interventions on DNAm biomarkers where each point is one intervention study. Significant effect sizes are marked with red asterisks (* p < 0.05, ** p < 0.01, *** p < 0.001), while solid lines indicate the mean effect size. F-G) Significant change count and effect sizes for pharmacological interventions, plotted analogously to the lifestyle interventions in D-E.

### Reliable Gen 2+ biomarkers align most with other biomarkers

When selecting epigenetic clocks to evaluate intervention responses, one may want to utilize a clock that is likely to agree with other clocks on the intervention effect. If the clock were to disagree with others, then the result would be confusing to interpret, or there would be a concern that the effect was a sporadic significant result (false positive). Thus we quantified the concordance of the 16 DNAm biomarkers - the likelihood that if a DNAm biomarker detected a significant effect then others would agree on the effect - analyzed across all interventions. Thus for each intervention where at least one biomarker responded significantly, we compared each significant biomarker with the other significant biomarkers in that study. If the biomarker agreed with greater than 50% of the remaining biomarkers we categorized this as an agreement, if less than 50% then a disagreement and if it was the only significant biomarker then as a third category of being “only significant”. Finally, for each of the 16 biomarkers, we divided the number in each category by the total number of significant results for that clock. We observed two trends (Figure 4C). First, Gen 1 biomarkers had less agreement with other biomarkers (Horvath1 = 0.66, Horvath 2 = 0.5, Hannum = 0.55) as compared to their Gen 2+ counterparts (DNAmEMRAge = 0.75, OMICmAge = 0.75, GrimAgeV1 = 0.63, PhenoAge = 0.83 and DunedinPoAm38 = 0.77). Second, reliable versions showed greater agreement compared to their original versions (Horvath2 0.5 vs PCHorvath2 0.77; Hannum 0.55 vs PCHannum 0.9; GrimAge 0.6 vs PCGrimAge 0.9; PhenoAge 0.83 vs PCPhenoAge 1; and DunedinPoAm38 0.76 vs DunedinPACE 0.82) with one exception (Horvath1 0.66 vs PCHorvath1 0.44). Of particular interest were cases where a given clock was the only one that showed any significant change. Most biomarkers that showed at least one result where they were the “only significant” clock also had many other results where they had large disagreements with other clocks. We interpret these clocks as having a high likelihood of false positives. However, there is one notable exception: DunedinPACE, which was unique in that it never disagreed with other clocks, yet it was the only significant clock in 17.6% of the interventions where it was significant. While it is still possible these “only significant” clock results for DunedinPACE are false positives, it is also possible that DunedinPACE is simply the most sensitive clock to intervention effects or can detect unique intervention effects.

### Different DNAm biomarkers are sensitive to different intervention categories

We further wanted to test whether different DNAm biomarkers are responsive to different categories of interventions. To test this, we replicated our analysis from Figure 4 A and B but this time limited to Lifestyle and Pharmacological interventions analyzed separately. In Lifestyle interventions (Fig 4D) DunedinPACE reported significant decreases in 8 out of the 15 interventions followed by SystemsAge (6), DunedinPoAm38 (5), DNAmEMRAge (4), and PCGrimAge (3). One-way t-tests revealed, (Fig 4E) that DunedinPoAm38 reported the largest decrease in effect size (mean = -0.1048, p = 0.0001), followed by DunedinPACE (-0.08887, 0.0037), Hannum (-0.08377, 0.015), and GrimAgeV2 (-0.07904, 0.0004). Interestingly, Horvath2 showed a significantly increased effect size (mean = 0.0802, p = 0.0054) and was the only clock to show an increase in one lifestyle intervention.

In comparison, after Pharmacological interventions (Fig 4F), GrimAgeV2 reported the highest number of significant decreases (8 out of 14) followed by PCGrimAge and OMICmAge (6). The top 4 biomarkers with largest decrease in effect size in the one way t-test analysis were DunedinPoAm38 (mean = -0.2394, p= 0.0442), GrimAgeV1 (-0.2332, 0.0151), GrimAgeV2 (-0.2092, 0.0058) and PCGrimAge (-0.1575, 0.0095). Although some biomarkers did overlap between the top categories, many biomarkers were sensitive to the intervention type and their magnitude of effect size differed between the two interventional categories. Most notably, DunedinPACE was altered by fewer Pharmacological interventions than other 2nd generation reliable biomarkers, despite its high responsiveness in Lifestyle (Figure 4D) and in uncategorized interventions (Figure 4A). Meanwhile, GrimAgeV2 was the most sensitive to Pharmacological interventions, but only changed in a single Lifestyle intervention. Thus, different biomarkers were most sensitive to different intervention categories.

### Study population health status is critical to DNAm biomarker response

Given that different interventional studies and trials in our analysis had clinical covariates and study metadata, we asked which of these may be affecting DNAm biomarker responsiveness. For this analysis, we built simple linear regression models associating the effect sizes from the different DNAm biomarkers with 8 different clinical covariates: Study population (Healthy vs Disease), number of subjects, intervention period, mean age, standard deviation of age, minimum age, age range, and percentage of females in the study (Fig 5A). Study population (Healthy vs Disease) was most strongly associated with highly responsive biomarkers (PCPhenoAge Z-score = 4.83; PCGrimAge = 4.07; SystemsAge = 4.65). This prompted the question: are DNAm biomarkers differentially responsive in Healthy and Disease population studies? We performed a one way t-test on effect sizes from interventions in Healthy groups (Fig 5B) and Disease groups (Fig 5C). The one-way t-test revealed multiple Gen 2+ reliable biomarkers (PCPhenoAge mean = -0.3203, p value = 0.0057; PCGrimAge m=-0.2226, p=0.0022; SystemsAge = -0.213, p = 0.0105, GrimAgeV2 m = -0.1874, p = 0.002;

**Figure 5:**
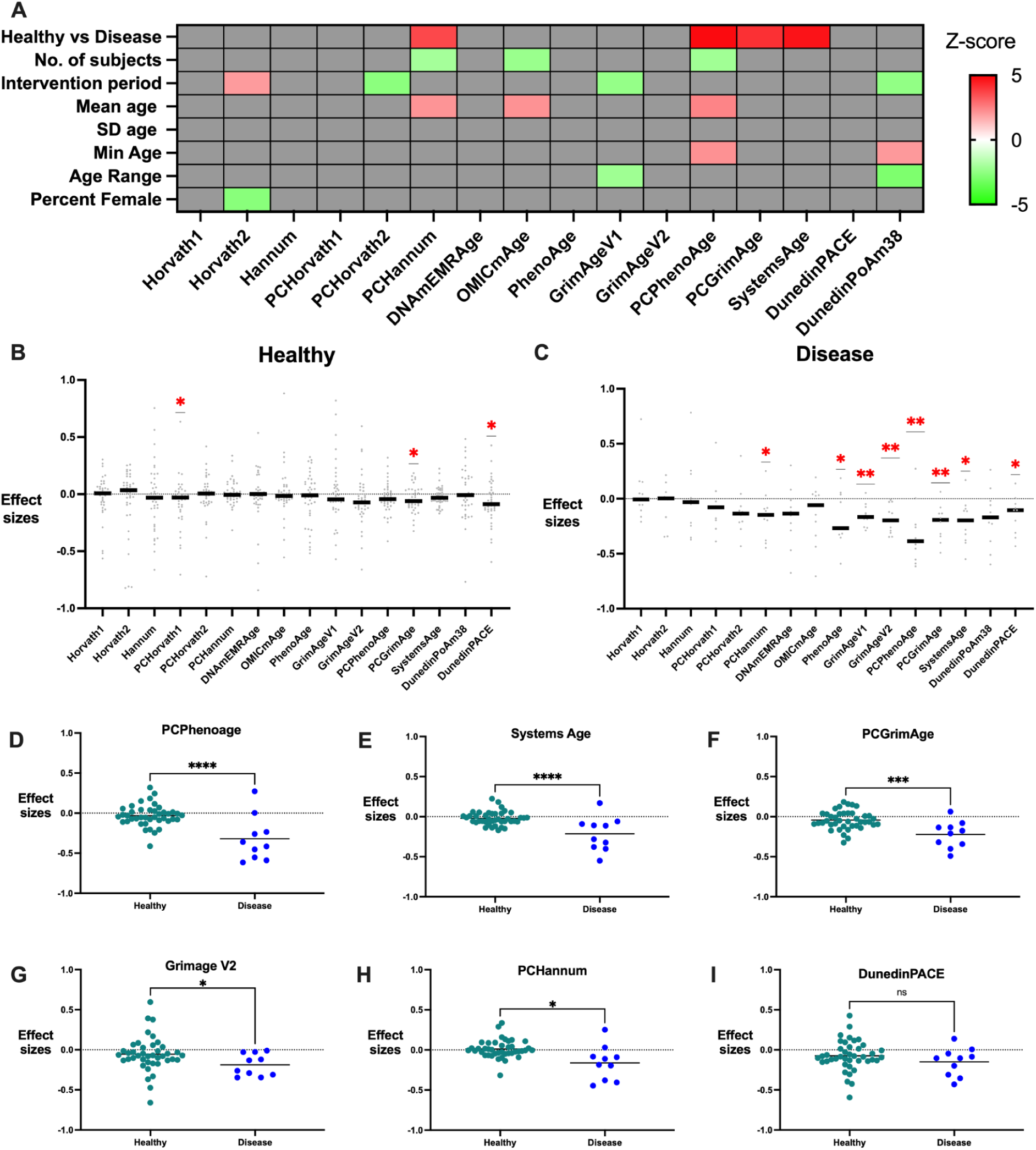
Effect of study characteristics on DNAm biomarker response to interventions. A) Heat map displaying the Z-scores of association from a linear model associating various study characteristics to DNAm biomarker effect sizes conducted across all interventions . Red signifies that the study characteristic results in higher effect sizes green indicates that the study characteristic results in lower effect sizes, and grey reflects no significant association. For Healthy vs. Disease, Healthy is the reference group so red indicates that studies in Disease populations show higher effect sizes. B-C) categorical scatter plots illustrating the effect sizes of DNAm biomarkers for studies conducted in healthy individuals (B) and individuals with diseases (C) Significant effects are marked with red asterisks.(* p < 0.05, ** p < 0.01, *** p < 0.001) D-I) Categorical scatter plots comparing effect sizes between healthy and disease subjects for specific biomarkers: D: PCPhenoAge, E: SystemsAge, F: PCGrimAge, G: GrimAgeV2, H: PCHannum, I: DunedinPACE.

DunedinPACE m= -0.1492, p = 0.0246), as well as PhenoAge (m = -0.1873, p = 0.035 ), GrimAgeV1 (m = -0.1529, p = 0.0015) and PCHannum (m = -0.1615, p = 0.0406) that responded to interventions with disease populations. On the other hand, only 3 DNAm biomarkers were responsive in Healthy populations: PCHorvath1 (mean = -0.06574, p-value = 0.0491), PCGrimAge (mean = -0.04327, p-value = 0.0191) and DunedinPACE (mean = -0.07445, p- value = 0.0143).

This led us to test whether the observed responsiveness of DNAm biomarkers was primarily driven by a handful of interventional studies in disease populations (10 studies) and not the majority of studies in healthy populations (41 studies). To test this, we performed an unpaired t-test on the distribution of effect sizes in Healthy versus Disease populations and found that many of the most responsive biomarkers were significantly more responsive in disease populations (Fig 5 D - I). PCPhenoAge (mean difference = 0.28, p-value = <0.0001), SystemsAge (0.2137,<0.0001) and GrimAgeV2 (0.139, 0.02) all had significantly greater decreases in disease populations as opposed to the Healthy populations. This was expected, as these biomarkers were responsive in the intervention studies in disease populations in Fig 5C and not in the healthy populations in Fig 5B. However, even PCGrimAge (mean 0.1793, p-value = 0.002) which was responsive in both Healthy and disease populations had a significantly greater decrease in disease populations as opposed to the Healthy populations. Thus, many biomarkers are responsive in Disease study groups but few are responsive in Healthy populations. The only DNAm biomarker which seems to be changing unbiasedly in both potentially healthy and disease populations seems to be DunedinPACE (mean difference = 0.07, p-value = 0.25) which was significantly different in both Healthy and Disease populations, but did not have a significant difference in effect sizes between the two populations.

### Gen Explainable DNAm biomarkers offer mechanistic insights into responsiveness

Clocks like SystemsAge, GrimAge, and OMICmAge were built from component DNAm biomarkers to better explain the specific aging changes that they are capturing. Thus, we also tested responsiveness for 78 Gen Explainable (GenX) DNAm biomarkers. We observed 39 biomarkers with significant changes across all interventions (Fig 6A). Examples of biomarkers with particularly high responsiveness included PCGrimAge components PCPACKYRS effect size = -0.05824(t=4.636), PCCystatinC -0.1404, 3.630), and PCTIMP1 (-0.08526, 3.666);

**Figure 6:**
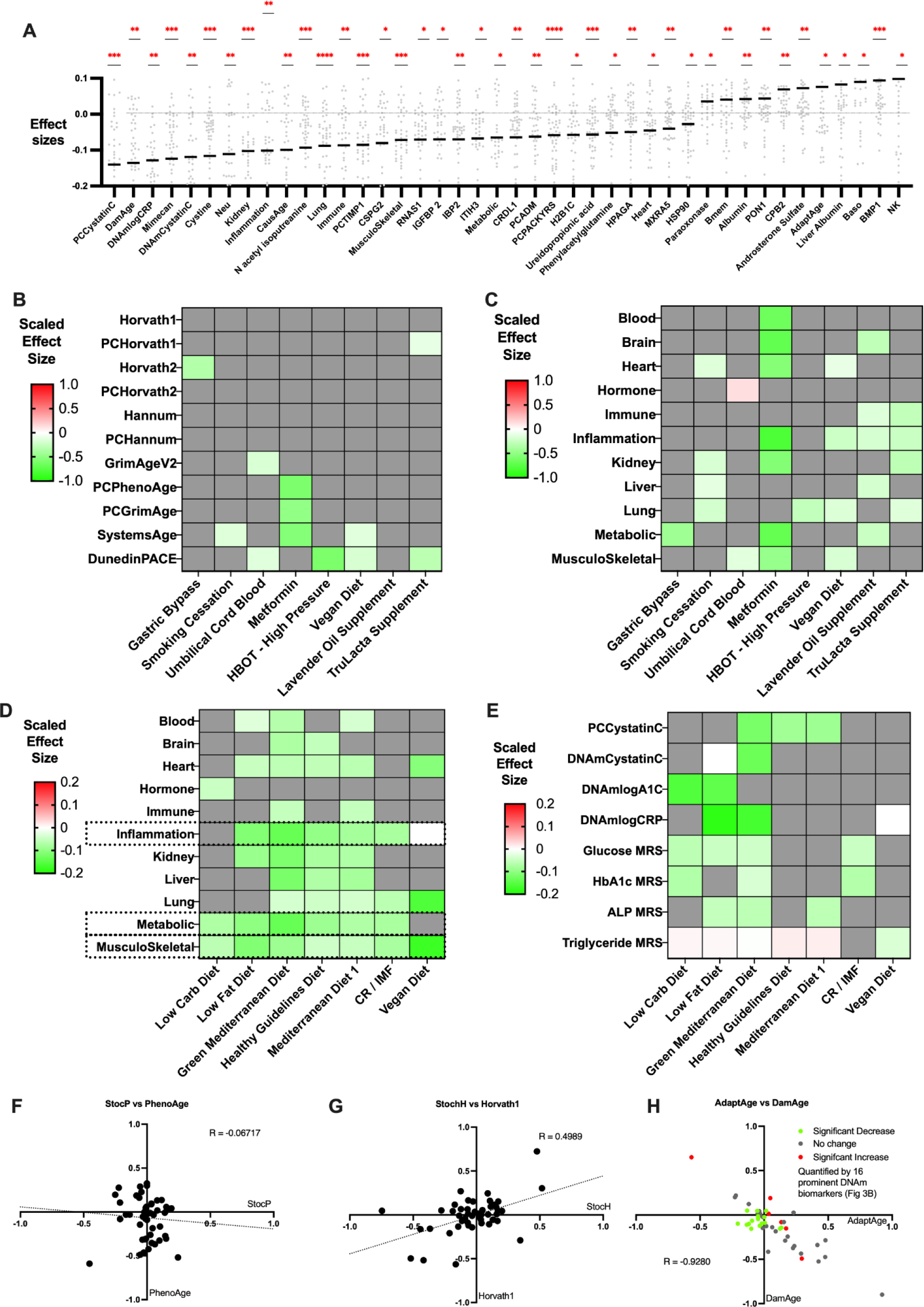
Explainable DNA methylation biomarkers (Gen X) show specific changes and provide mechanistic insight. A) Scatter plot of effect sizes (Y axis) for significant Gen X DNAm biomarkers (X axis). Significant effects are marked with red asterisks.(* p < 0.05, ** p < 0.01, *** p < 0.001) B) Heatmap of whole body aging DNAm biomarkers (Y axis) and specific interventions (X axis). Red indicates increase in epigenetic age, green indicates decrease in epigenetic age, while gray denotes no significant effect. C) Heatmap of system specific DNAm biomarkers (Y axis) and specific interventions (X axis). The color scale is consistent with Panel B. D) Heatmap of Scaled Effect Sizes for Different Organ Systems and Diets. The color schema is consistent Panels B and C. E) Heatmap of Scaled Effect Sizes of specific metabolite and protein DNAm proxies and Diets. The color scale is consistent with Panels B-D. F) Scatter Plot of StocP (X axis) vs. PhenoAge (Y axis) effect sizes. G) Scatter Plot of StocH (X axis) vs. Horvath1 (Y axis). H) Scatter Plot of AdaptAge (X axis) vs. DamAge (Y axis). Red represents interventions that significantly increase epigenetic age across 16 key aging biomarkers, green indicates those that significantly decrease epigenetic age, and gray shows no significant impact on epigenetic age across the same 16 biomarkers.

SystemsAge components Kidney (-0.1021, 3.753), Lung (-0.08812, 4.615), and MusculoSkeletal (-0.07132, 3.729); and finally OMICmAge components EBP-N-acetyl-isoputreanine (-0.09316, 3.506), EBP-Ureidopropionic acid (-0.05658, 3.733), EBP-Mimecan (-0.1238, 4.047), EBP-Cystine (-0.1162, 3.897), EBP-CBPB2 (0.06373, 3.497) and EBP-BMP1 (0.09398, 4.183).

To understand how GenX clocks can provide more insight into responsiveness, we examined specific intervention, selected based on whether they fell into two categories: (1) significantly decreasing epigenetic age in only a handful of Gen2 reliable clocks (Smoking Cessation - SystemsAge; UCB - DunedinPACE and GrimAgeV2; Metformin - SystemsAge, PCGrimAge, PCPhenoAge; HBOTHP - DunedinPACE; Vegan Diet - DunedinPACE and SystemsAge; TruLacta Supplement - DunedinPACE) or (2) not significantly decreasing epigenetic age in any Gen2 reliable clocks (Gastric Bypass, Lavender Oil Supplement LOS) (Figure 6B). The sporadic responsiveness of clocks to the first category of interventions may suggest they are false positives, while the lack of responsiveness in the second cateogry may suggest they do not modify epigenetic age. However, we considered the possibility that these interventions simply had very specific effects that may not be detectable using a general aging clock. Accordingly, the SystemsAge components suggested these interventions may be impacting different systems (Figure 6C). For example, the two interventions that did not modify any general Gen 2 reliable clock do show significant decreases in epigenetic age of specific systems (Gastric Bypass decreases Metabolic score with effect size 0.43; LOS decreased Brain 0.29, Immune 0.15, Inflammation 0.19, Kidney 0.2 and Metabolic 0.24). Other interventions Smoking cessation seeing maximal decrease in Lung (0.20), UCB in MusculoSkeletal (0.13), and HBOTHP in Lung (0.27). While other systems saw multi-system decreases like Metformin with Inflammation (0.80), Brain (0.70), and Metabolic (0.70) decreasing the most. In the Vegan diet, the largest decrease was seen in Inflammation (0.23) and Musculoskeletal (0.17). While in TruLacta supplement, Kidney (0.29), Immune (0.28) and Inflammation (0.24) reported the greatest impact after the intervention.

Finally, given we had a multitude of diets (7) in our study (Low Carb Diet, Low Fat diet, Green Mediterranean Diet, Healthy Guidelines Diet, Red Meat Mediterranean Diet, CR/IMF, Vegan Diet) we wished to understand if GenX DNAm biomarkers could provide deeper mechanistic understanding specifically into dietary interventions. We looked at the 11 system scores and specific metabolic related protein and metabolite epigenetic proxy biomarkers (Figure 6 D and E). Among the 11 system specific biomarkers, Musculoskeletal was the only score that significantly decreased across all 7 diets. Meanwhile, two system scores (Metabolic and Inflammation) were responsive in 6 of the 7 diets. When looking at the protein and metabolite proxies, we observed the epigenetic proxy of Triglycerides to be significant across 6 of the 7 diets while the glucose proxy was significant in 4 of 7 diets, suggesting great potential for these epigenetic proxies in measuring diet-based interventions.

### Responsiveness of stochastic and causal components

A new generation of explainable DNAm biomarkers suggest they have both stochastic and non-stochastic components ^32^, as well as causal components^18^ that may be either damaging or adaptive in the aging process. It has been shown that Gen1 DNAm biomarkers that predict chronological age preferentially capture stochastic epigenetic noise, while Gen2+ DNAm biomarkers capture a larger proportion of non-stochastic aging processes.^32^ However, this analysis was done on cross-sectional data, so we aimed to leverage TranslAGE-Response to test this using many longitudinal datasets.

We correlated the effect sizes of the stochastic component of Horvath 1 (StocH) with the full Horvath1 clock, and did the same for PhenoAge (StocP vs PhenoAge) (Figure 6 F-G). We observed PhenoAge had little correlation with StocP (R=-0.06) while Horvath1 had high correlation with StocH (R= 0.50). Thus, our longitudinal data support the notion that changes captured by Gen1 DNAm biomarkers such as Horvath1 are of the accumulation of stochastic epigenetic noise, while those changes captured by Gen2+ DNAm biomarkers are more non-stochastic.

It was previously shown in three datasets that aging interventions decrease damaging causal components (DamAge), while one of those also showed an increase in adaptive causal components (AdaptAge). We used translAGE-Response to verify that there is indeed a strong inverse relationship between DamAge and AdaptAge changes (R=-0.8929) after interventions. We also asked whether the interventions that tend to decrease the 16 biomarkers in Fig. 3B tend to have negative DamAge but positive AdaptAge, while those that increase those 16 biomarkers tend to have negative AdaptAge but positive DamAge. We did not observe this trend, rather most of the significant interventions were in Q3 where AdaptAge and DamAge were both negative (Figure 6H).

## Discussion

Epigenetic clocks, or more generally DNAm aging biomarkers, have long been prophesied as measures that could become surrogate endpoints in longevity intervention clinical trials. However, these prophecies have been based on their prognostic capabilities, that is, their ability to associate with future mortality and morbidity. Various optimizations to these DNAm biomarkers have recently been proposed that would make them useful for clinical trials, such as increasing test-retest reliability and longitudinal performance, increasing their explainability , enriching in causal changes, and reducing their vulnerability to confounding variables. However, what has never been tested in a systematic manner is the responsive nature of these biomarkers, comparing across many biomarkers and many interventions simultaneously. Thus, in this publication, we set out to answer the question “Are DNAm aging biomarkers responsive?".

To do this, we curated TranslAGE-Response, a database of 51 longitudinal longevity intervention studies from public (GEO, EMBL) and private databases, selecting those with whole blood DNA methylation data (450K or EPIC) pre- and post-intervention. Metadata was standardized, and 110 DNAm aging biomarkers were calculated and scaled to the Health and Retirement Study for comparability. Age-residualized scores were computed and analyzed using paired t-tests to assess pre- and post-intervention differences. Effect sizes were visualized in a heatmap to explore which DNAm biomarkers and interventions showed the strongest responses and factors influencing biomarker responsiveness.

Our data revealed that pharmacological interventions induced the strongest responses in DNAm biomarkers compared to lifestyle changes, supplements, and medical procedures. Pharmacological agents, such as anti-TNF therapies and metformin, consistently modified DNAm biomarkers, indicating their robust impact on biological aging processes. The substantial effects observed with these pharmacological interventions suggest their potential as frontline options in longevity therapies. The observed effects of pharmacological interventions may be due to their direct influence on molecular pathways that regulate aging and inflammation. For example, anti-TNF therapies can reduce systemic inflammation^33^, a key driver of biological aging, thereby slowing down the aging process at the molecular level and has been shown to extend lifespan in mice while having additional health benefits.^34–36^ Similarly, metformin, known for its effects on metabolic pathways, may enhance cellular resilience and longevity by modulating pathways such as AMPK and mTOR. Future research should delve deeper into the specific molecular pathways affected by these interventions to fully understand their mechanisms of action and optimize their use in clinical practice.

The study also evaluated the replicability of interventions in modifying DNAm biomarkers. We found that adherence to the principles of consistency—modifying the same biomarkers to similar extents within (Rule 1) and across studies (Rule 2) —enhances the reliability of the results of an intervention. Interventions such as anti-TNF therapies and the Mediterranean diet demonstrated consistent modifications, underscoring their robustness as longevity interventions while Senolytics and SRW supplement did not. The reproducibility of these findings across multiple studies emphasizes the importance of standardized methodologies in the assessment of interventions.

Our results also highlighted that advanced DNAm biomarkers, particularly those from Generation 2+ (Reliable and mortality or rate of aging predictors) such as GrimAgeV2, PCGrimAge, PCPhenoAge, SystemAge and DunedinPACE, exhibit substantial responsiveness to these interventions. Specifically, these biomarkers showed profound changes in individuals undergoing longevity treatments, reinforcing their potential utility in both research and clinical settings for monitoring the effectiveness of anti-aging strategies. The discriminative power of these advanced biomarkers makes them essential tools in personalized medicine for tracking biological age and intervention efficacy.

A key observation from our analysis was the variability in responsiveness among DNAm biomarkers, influenced significantly by the type of intervention. For instance, while PCPhenoAge showed variable results across different studies, GrimAgeV2, SystemsAge, PCGrimAge and DunedinPACE consistently indicated significant decreases in epigenetic age, regardless of the intervention type (Lifestyle or Pharmacological). This consistency highlights the crucial role of selecting suitable biomarkers for specific interventions, ensuring that measurements of biological aging remain reliable and accurate. These findings suggest that reliable Gen 2+ DNAm biomarkers should be prioritized in clinical trials to provide robust insights.

Another critical factor influencing biomarker responsiveness was the health status of the study population. Our study found that DNAm biomarkers were significantly more responsive in disease populations compared to healthy populations. For example, biomarkers like PCPhenoAge, SystemsAge, and GrimAgeV2 showed greater decreases in epigenetic age in disease individuals than their healthy counterparts, suggesting that these populations might benefit more distinctly from anti-aging interventions. This finding is crucial as it implies that individuals with existing health conditions stand to gain the most from targeted anti-aging interventions. The heightened responsiveness in disease populations may be attributed to the underlying pathological processes that accelerate biological aging, making these individuals more sensitive to the effects of interventions aimed at slowing or reversing these processes. This insight highlights the necessity of stratified clinical trial designs that consider the baseline health status of participants. Tailoring interventions to specific health profiles could optimize the therapeutic outcomes and enhance the generalizability of findings across diverse populations.

We also saw that explainable or Gen X DNAm biomarkers provided deeper mechanistic insights into the effects of specific interventions. By examining system-specific scores, we identified targeted effects on different physiological systems. For instance, smoking cessation primarily decreased lung-specific system score, whereas metformin exhibited extensive effects across inflammatory, brain, and metabolic systems. These findings suggest that Gen X biomarkers can uncover nuanced responses to interventions that whole-body clocks might overlook. Understanding the specific systems affected by various interventions can inform the development of targeted therapies that address the underlying causes of aging in specific organs or tissues. For example, interventions that target inflammation and oxidative stress in the vascular system can help reduce cardiovascular aging, while those that enhance neuroplasticity and reduce neuroinflammation can support healthy brain aging. The ability to dissect the specific systems impacted by various interventions enhances our understanding of the underlying biology of aging and the mechanisms of action of anti-aging therapies. Additionally, leveraging systems biology approaches and integrative omics technologies can further elucidate the complex interactions between different biological systems and how they contribute to the aging process.

In conclusion, our study establishes a foundational framework for understanding the responsiveness of DNAm biomarkers to various longevity interventions. Generation 2+ reliable biomarkers, particularly DunedinPACE, emerged as leading indicators of biological age response. The differential responsiveness across intervention types and population health statuses emphasizes the necessity of tailored biomarker selection in clinical trials. Future research should prioritize validating these biomarkers as surrogate endpoints to advance the development of effective anti-aging therapies. We hope this work encourages “clock-makers” and “DNAm biomarker-builders” to not only evaluate the prognostic value of their biomarkers but also prioritize assessing their responsiveness. By making responsiveness a key step in the validation process, scientists can advance the development of DNAm biomarkers that are both predictive of aging and adaptable to longevity interventions, ultimately improving their utility in clinical and research settings. We also anticipate that this work will motivate future randomized clinical trials aimed at validating both the responsiveness and the surrogate endpoint status of DNA methylation biomarkers and eventually translating these findings into practical applications that improve health outcomes and quality of life for individuals across the lifespan.

## Acknowledgments

This work was supported by the National Institute on Aging (NIA:1R01AG065403 to A.H.C.). It was also supported by the Impetus Grant (R.S.), the Gruber Science Fellowship at Yale University (R.S.), and the Thomas P. Detre Fellowship Award in Translational Neuroscience Research from Yale University (to A.H.C.).

## Author Contributions

R.Sehgal, D.S.B. and A.H.C. conceived the project and study design. R.S. performed responsiveness analysis. R.Sehgal, D.S.B., J.K., J.A., and A.H.C. built the pipeline for TranslAGE. R.Sehgal, D.S.B. J.K, J.A., Y.M., J.G., A.P., A.H.C, V.D. and N.C. performed study data and metadata curation. A.H.C. and R.Sehgal provided supervision and managed the overall project. All authors reviewed and contributed to the manuscript.

DNAm aging biomarkers community - provided clock code

Longevity interventional studies community - provided datasets for longevity interventional analysis.

## Data Availability Statement

All effects sizes will be posted upon publication. A separate publication is in preparation on a platform to share the publicly available subset of datasets and potentially others in a cleaned and harmonized format. Code to calculate all clocks except for OMICmAge and DNAmEMRAge will be accessible at https://github.com/HigginsChenLab/methylCIPHER after publication. Code to calculate OMICmAge, DNAmEMRAge, and underlying algorithms will be accessible via TruDiagnostic’s DNAm Analysis Software after publication. You can request access to the software at https://www.trudiagnostic.com/softwarerequest."

## TranslAGE availability

We intend for TranslAGE to become available as a resource to the geroscience community. If you are interested in utilizing translAGE for academic or commercial purposes, please let us know your ideas for translAGE applications. For more details, please

## Methods

### Analysis Pipeline

#### Step 1: Data Curation

We began by compiling a comprehensive list of publicly and privately available clinical trials and studies focused on longevity interventions. Publicly available datasets were sourced from Gene Expression Omnibus (GEO) and the European Molecular Biology Laboratory (EMBL) repositories, while private datasets were obtained from TruDiagnostic. Datasets were selected on the basis of the following criterion: 1) Interventional study or trial is of a intervention that has been hypothesized for the purposes of longevity or healthspan extension as defined in literature, A) 2) DNAm data is available pre and post the intervention, 3) Tissue of origin for the DNAm data is blood (ideally PBMCs), 4) Study passes quality checks such as: minimum number of individuals in the treatment arm and consistent methodology of DNAm data generation at pre and post time points. For each study, we curated extensive study-level metadata, including the following key variables:

● **Intervention type**: Identifying whether the intervention involved lifestyle changes, pharmacological treatments, or other longevity-enhancing approaches.
● **Diseases in population**: Listing any comorbidities or conditions present in the study population that could influence the effects of the intervention.
● **Age range and percent female**: Capturing demographic distribution to understand age- and sex-related response variability.

These metadata were manually extracted from study documentation and supplementary materials to ensure accuracy.

#### Step 2: Metadata standardization

We utilized a standardized cleaning script that was adapted for each dataset. We standardized the variables across all studies to create uniform, comparable datasets. This harmonization process involved:

● **Standardized columns**: We ensured that all datasets contained consistent variables, including **Age**, **Sex**, **Sample ID**, **Individual ID**, **Follow-up time from baseline draw**, and **Sample type** (control vs. subject). This step allowed for consistent cross-study analysis.
● **Sample identification**: To maintain data integrity, each sample was assigned a unique **Sample ID** and linked to an **Individual ID**, ensuring that repeated measurements from the same individual could be tracked longitudinally.

This harmonization facilitated subsequent analyses, allowing for robust comparisons between different studies and intervention types.

#### Step 3: Clock Calculation

For each of the curated datasets, we calculated over **110 DNA methylation (DNAm) biomarkers** using the **MethylCIPHER 2.0** package. This package allows for the computation of various methylation-based aging and health biomarkers, such as:

● **Epigenetic clocks** (e.g., Horvath, Hannum clock)
● **Health-related markers** (e.g., DNAmPackYears)
● **Protein and Metabolite Proxies** (e.g., DNAmGDF15, MRS Triglyceride)

These biomarkers were calculated for each sample based on the methylation data available in each study. A list of all the biomarkers calculated with additional information on them can be found in Supplementary Table 1.

#### Step 4: Age-residualized scores

We processed the biomarkers by age-residualizing them in each dataset. Age-residualization was performed to control for the natural variation in DNAm biomarkers due to chronological age. The procedure involved linear regression. We used age as the independent variable and each DNAm biomarker as the dependent variable to calculate the residuals. These age-residuals represent the portion of the biomarker that is not explained by chronological age, allowing us to isolate the effect of the intervention on DNAm biomarkers.

#### Step 5: Biomarker standardization

To ensure comparability across datasets, we standardized the calculated biomarkers with respect to the **standard deviation in the Health and Retirement Study (HRS)**. This standardization process involved calculating the standard deviation in HRS for all age residualized DNAm biomarkers. Since post linear regression means for all the age residuals for biomarker are already 0, all we needed to do was divide the age residuals for each DNAm biomarker by the standard deviation calculated in HRS for that biomarker to get them all at the same scale and comparable to each other.

#### Step 6: Paired t-test

Since all our studies had both **pre- and post-intervention samples**, we performed **paired t-tests** to evaluate the effect of the intervention on the calculated DNAm biomarkers. This analysis was conducted as follows:

● **Pre- vs. post-intervention comparison**: For each individual, the change in age residualized biomarker values between pre- and post-intervention samples was calculated. The paired t-test was used to assess whether these changes were statistically significant.
● **Significance threshold**: A p-value of <0.05 was considered statistically significant.

#### Step 7: Heatmap and follow-up analysis

To further explore the responsiveness of DNAm biomarkers to longevity interventions, we generated **heatmaps** and performed additional statistical analyses:

● **One-way t-tests**: Conducted to test for deviations from zero effect size in intervention responses.
● **Exploratory analyses**: Multiple analyses, including correlation tests and regression models, were employed to explore relationships between biomarker changes and other variables such as age, sex, and intervention type.

### One-way T-test testing for deviations from 0 effect size

To evaluate whether the observed effect size significantly deviated from zero, a one-sample t-test was performed using GraphPad Prism (Version 9.4.0). The dataset included different measurements based on the condition being tested (for example: 51 for all interventions, 41 for healthy, 10 for disease).Test Setup: In Prism, a one-sample t-test was conducted by specifying a hypothetical mean (Mu) of 0, which represents the expected effect size under the null hypothesis of no effect.: Normality of the data was assumed, and Prism automatically performed the test using the standard t-test formula. Prism provided the t-statistic, degrees of freedom (df), and two-tailed p-value to evaluate whether the mean effect size significantly differed from zero. A significance level of 0.05 was used. All analyses and visualizations were conducted within Prism.

### Clinical trial covariates association analysis

To assess the association between DNA methylation (DNAm) biomarkers and clinical covariates, we first curated a comprehensive set of clinical variables from the study datasets. Clinical covariates were selected based on their relevance to clinical trials. The covariates were extracted from study metadata or clinical records provided within each dataset. Where applicable, continuous variables were normalized or transformed to achieve better model fit. The association between DNAm biomarker effect sizes and clinical trial covariates was assessed using univariate regression models. This analysis aimed to identify covariates that significantly influenced DNAm biomarker responsiveness. For each DNAm biomarker, univariate linear regression models were fit to evaluate the independent association with each clinical covariate:

Biomarker = β0 + β1×Covariate + ɛ

Covariates were tested individually for each biomarker, and the strength of association was assessed using Z-score and absolute Z-scores less than 1.96 were considered not significant.

